# Systematic Scale-Up and Enhanced Purification of Marine Cyanophage P-SSP7

**DOI:** 10.64898/2026.01.03.697218

**Authors:** Pavlo Bohyutski, Amar D. Parvate, Natalie C. Sadler, William B. Chrisler, Margaret S. Cheung, James E. Evans

## Abstract

Cyanophages represent important models for understanding virus-host interactions, yet high-resolution structural studies remain relatively few due to challenges with preparing enough sample of sufficient quality for cryo-EM and functional multi-omics studies. Here we developed an integrated methodology for scaling production of the model cyanophage P-SSP7 from laboratory maintenance volumes (5-100 mL) to production scales (up to 40 L) while dramatically improving the quality of phage preparation for structural applications. Our systematic approach integrates host cultivation using adaptation to local seawater to reduce production costs, optimized infection protocols to maximize infectious titer yields, and multi-stage purification workflows specifically designed for cryo-EM quality requirements. The final methodology consistently produces infectious phage titers exceeding 3×10^12^ units/mL with recoverable yields of 10^13^ total infectious units and >95% purity validated by cryo-electron microscopy at each optimization step. Most significantly, this approach achieves a 60-fold reduction in cryo-EM data collection time by increasing usable particles per field of view for single particle analysis. Overall, our final preparations demonstrate robust phage stability, retaining 68% infectivity after 3 months and 23% after 6 months at 4°C. This workflow moves cyanophage culturing and downstream structural studies from specialized, resource-intensive endeavors toward routine research capability and establishes an adaptable framework for scaling production that can be applied to other host-virus systems.

## 1. Introduction

Picocyanobacteria are the most abundant photosynthetic organisms on Earth (Flombaum et al., 2013) and cyanophages that infect *Prochlorococcus* and *Synechococcus* are among the most abundant biological entities on the planet. These viruses control cyanobacterial populations, redirect metabolic fluxes, influence biogeochemical cycles, and drive microbial evolution through horizontal gene transfer (Rodriguez-Valera et al., 2009; Thompson et al., 2011; Rozum et al., 2025). Understanding these virus-host interactions at the molecular level requires high-resolution structural details that can be revealed by techniques such as cryo-electron microscopy and electron tomography to resolve capsid architecture, surface protein organization, and genome packaging mechanisms, and to track time-resolved dynamic infection processes (Milne et al., 2012; Bai et al., 2015; Wrapp et al., 2020). However, such studies on cyanophages have been severely constrained by methodological challenges in producing high-quality, concentrated viral preparations.

Specifically, high-resolution cryo-EM requires maximizing intact, usable particles per microscope field to minimize images needed and conserve instrument time. This drives requirements for very high particle concentrations of up to 10¹¹-10¹² particles/mL. Additionally, particles must be highly purified to enable effective computational processing during 3D reconstruction, as contaminating debris or broken/damaged phages can interfere with particle picking algorithms and complicate downstream structural refinements. Achieving these requirements for cyanophages infecting *Prochlorococcus* presents particular challenges. First, the fastidious growth requirements of cyanobacterial hosts demand specialized media, precise environmental control, and extended cultivation periods (Moore et al., 2007; Morris et al., 2011).

While commercial Sargasso Seawater can be used as the base for *Prochlorococcus*-specific media, it costs $85 per L plus shipping (NCMA, 2025), so it can become prohibitively expensive at production scales. Second, cyanophage production itself is highly sensitive to host cultivation and infection conditions, with factors such as light intensity, temperature, pH, multiplicity of infection (MOI), carbon dioxide, and nutrient availability dramatically affecting phage synthesis kinetics, yield, and the frequency of defective particles that compromise sample quality (Wilson et al., 1996; Cheng et al., 2017; Cheng et al., 2019; Laurenceau et al., 2021; Yadav and Ahn, 2021; Zhao et al., 2022; Jo et al., 2024). Third, cyanobacterial lysis generates complex lysates containing cellular debris, membrane vesicles, lipopolysaccharides, and empty capsids that require sophisticated multi-stage purification to achieve cryo-EM-grade purity (Wolf and Reichl, 2014; Tanir et al., 2021).

While recent hardware advances in cryo-EM instrumentation and image processing software have enabled protein structure determination at resolutions as high as ∼1.3 Å (Nakane et al., 2020; Yip et al., 2020), sample preparation has emerged as the primary constraint limiting structural studies (Passmore and Russo, 2016; Carragher et al., 2019; Weissenberger et al., 2021). Studies show that even under optimal conditions, substantial fractions of cryo-EM exposures fail to meet quality standards required for high-resolution reconstruction, with some reports indicating failure rates of ∼50% (Alewijnse et al., 2017; Clare et al., 2017). This challenge becomes particularly acute for marine viruses given their high salt requirements in media and the additional complexity of achieving adequate particle concentrations while maintaining structural integrity. The combination of these production bottlenecks coupled with generally limited cryo-EM access creates a compounding problem where valuable microscope time could be wasted on suboptimal preparations that cannot support high-resolution structural determination. Cryo-EM instruments cost millions (USD) to purchase with yearly service contracts requiring hundreds of thousands (USD) in maintenance costs. Two access modes typically exist for these instruments: either an institution owns a cryo-EM and makes it available to staff at an hourly recharge rate, or regional or national centers exist that provide free access to users but may have multiple-week wait times to access the instrument between sessions. Therefore, to make structural studies on P-SSP7 and other cyanophages more feasible, an improved approach is needed to routinely generate high-yield and high-quality preparations of intact viruses.

Cyanophage P-SSP7 infects *Prochlorococcus marinus* strain MED4 and serves as an important model system for understanding virus-host interactions and population dynamics (Lindell et al., 2007; Murata et al., 2017; Schwartz and Lindell, 2017; Liu et al., 2019; Laurenceau et al., 2021; Rozum et al., 2025). P-SSP7 is among only a few cyanophages with structural information in the PDB (Liu et al., 2010; Cai et al., 2023) and belongs to a class of tailed phages which represent an exquisite assembly with proteins in the capsid, portal complex, and tails arranged in several different symmetries (Cai et al., 2023; Li et al., 2023; Sonani et al., 2024). However, the structural story for P-SSP7 is incomplete as the models in the PDB only provide coordinates for the P-SSP7 capsid architecture determined from an intermediate resolution reconstruction at 4.6 Å and lack details for the portal-tail complex, which is known to be crucial for infection (Liu et al., 2010; Rao et al., 2021; Cai et al., 2023). This limitation stems from the inherent symmetry mismatch between the icosahedral capsid (5-fold symmetry at vertices) and the portal-tail apparatus (12-fold portal, 6-fold tail) and is a shared trait of P-SSP7 and numerous other tailed bacteriophages (Hendrix, 1978; Li et al., 2022). Unfortunately, the issue of symmetry mismatch is a fundamental challenge which prevents the use of symmetry averaging and necessitates significantly larger imaging datasets for high-resolution asymmetric reconstruction (Morais et al., 2001; Huiskonen, 2018). Intriguingly, new symmetry maximization and marginalization routines in cryo-EM software like CryoSPARC (Punjani et al., 2017) and Relion (Burt et al., 2024) can overcome the issue of symmetry mismatch, making the production of high-quality samples the only remaining bottleneck for high-resolution complete structure determination of P-SSP7.

Current cyanophage production methods typically yield inadequate samples more suitable for basic characterization studies. While individual techniques for *Prochlorococcus* cultivation (Moore et al., 2007; Chisholm Laboratory, 2022) as well as collections of protocols for scaled phage production, concentration, and purification exist (Thurber et al., 2009; Hurwitz et al., 2012; Luong et al., 2020; João et al., 2021; Lapras et al., 2024; Luong et al., 2024), these workflows were either developed for specialized therapeutic and viral metagenome (virome) applications, or are not easily duplicated across laboratories due to insufficient detail, or remain fragmented and fail to address the integrated challenges of cost-effective scaling, infection optimization, and cryo-EM-grade purification. Most critically, existing methods do not account for the environmental sensitivity of cyanophage production—cultivation conditions can dramatically affect both phage infectivity and the ratio of intact to defective particles (Wilson et al., 1996; Cheng et al., 2017; Cheng et al., 2019; Laurenceau et al., 2021; Yadav and Ahn, 2021; Zhao et al., 2022)—nor do they achieve the >95% purity and 10¹¹-10¹² particles/mL concentrations essential for comprehensive structural studies. The absence of integrated workflows specifically designed for structural applications has created a fundamental bottleneck preventing the realization of cryo-EM’s theoretical capabilities for virus systems. For P-SSP7 in particular, we found that adapting protocols described in prior publications on partial structure determination by cryo-EM and visualization of initial infection dynamics with cryo-EM/ET to our laboratory context and scale-up goals required considerable assumptions and modifications to various steps. We therefore sought to create a more robust approach to the preparation of cyanophage samples for structural and multi-omics studies using P-SSP7 as the model phage.

Here, we address these limitations through a systematic approach for scaling P-SSP7 cyanophage production from 5 mL laboratory cultures to 40 L production volumes. Our methodology integrates cost-effective local seawater adaptation and optimized cultivation protocols for host *Prochlorococcus* with quality-driven infection and purification workflows. The approach consistently achieves infectious phage titers of 3.1×10¹² units/mL with total recoverable yields of 10¹³ infectious units and >95% purity, meeting the stringent requirements for high-resolution structural studies. Most significantly, this approach increases usable particle density from ∼1 particle per average image field of view to 66 particles per field, decreasing data collection time by a factor of 60× to yield the same number of final particles. Additionally, we show that the resulting purified phages also enable visualization of host-phage interactions through controlled infection experiments.

## 2. Materials and Methods

Our systematic scale-up methodology progressed through three phases, each addressing specific technical challenges while building toward production volumes and sample quality suitable for structural studies. The small-scale phase (5-100 mL) adapted existing protocols (Moore et al., 2007; Morris et al., 2011) to our laboratory conditions, establishing baseline performance metrics. The intermediate-scale phase (1-8 L) implemented cost-effective local seawater adaptation and custom cultivation systems with modified aeration protocols based on recommendations in (Moore et al., 2007; Chisholm Laboratory, 2022), along with initial optimization of infection procedures and purification methods, while cryo-EM validation revealed critical sample quality limitations. The large-scale phase (10-40 L) completed infection and phage harvesting optimizations while implementing enhanced purification workflows designed to meet cryo-EM particle density and purity requirements based on established viral purification methods (João et al., 2021; Lapras et al., 2024; Luong et al., 2024). Throughout all phases, cryo-EM imaging served as the primary quality control metric following best practices for sample assessment (Passmore and Russo, 2016; Carragher et al., 2019; Weissenberger et al., 2021). Particle density and debris assessments directly informed purification refinements and scale advancement decisions.

### 2.1. Host Culture Maintenance and Local Seawater Adaptation

#### Cyanobacterium Cultivation

*Prochlorococcus marinus* strain MED4 (CCMP1986) was obtained from the Provasoli-Guillard National Center for Marine Algae and Microbiota (NCMA, East Boothbay, ME) and maintained in Sargasso seawater-based Pro99 medium under established protocols and strictly controlled environmental conditions (Moore et al., 2007; Morris et al., 2011). Cultures were grown under a 12-h light:12-h dark cycle with light intensity of 25 μmol quanta m ²s ¹ and constant temperature of 21°C in temperature-controlled growth chambers. These specific cultivation conditions were selected based on optimized parameters for MED4 growth (Moore et al., 2007; Morris et al., 2011) and to ensure consistency with previous P-SSP7 structural studies (Liu et al., 2010; Murata et al., 2017). Small-volume cultures (5-10 mL) were maintained in Falcon® 5 mL Round Bottom Polystyrene Test Tubes with snap caps (#352054, Corning Incorporated, USA), while larger maintenance cultures (25-100 mL) utilized CELLSTAR® Filter Cap Cell Culture Flasks with 25 cm² or 75 cm² growth surface area (#690175 or #658175, Greiner Bio-One International GmbH, USA). Cultures were routinely transferred every 2-3 weeks during mid-exponential growth phase using a 5-10% inoculum to maintain log-phase growth characteristics and prevent culture senescence.

#### Local Seawater Adaptation Protocol

To enable cost-effective scale-up while maintaining culture viability, *P. marinus* MED4 cultures were systematically adapted from commercial Sargasso Seawater to local Salish Sea seawater over a two-month period. Commercial Sargasso Seawater presents significant cost and logistical constraints for large-scale phage production, while locally sourced seawater (*e.g.*, Salish Sea water collected at PNNL Sequim dock) offers a more economical alternative. However, these different seawater sources differ substantially in salinity: Sargasso surface waters measure 36-37.7 PSU (Not et al., 2007) compared to 29-20 PSU in Salish Sea surface waters (Khangaonkar et al., 2019; Sobocinski, 2021). While MED4 can adapt to salinities as low as 28 PSU (He et al., 2022), such adaptation to changes in medium composition should be performed gradually to allow cyanobacteria to deploy the salt-acclimation mechanisms they possess (Hagemann, 2011; Pade and Hagemann, 2014). This gradual adaptation approach can substantially reduce production costs while maintaining culture viability for large-scale applications.

The adaptation protocol involved incremental increases in local seawater proportion (0% → 20% → 40% → 60% → 80% → 100%) with 1-2 weeks of stabilization at each concentration. Culture density and growth rates were monitored using optical density measurements and flow cytometry to ensure successful acclimation before advancing to the next step. Local seawater was filtered through 0.2 μm filters and autoclaved at 121°C for 20 minutes before use to eliminate contaminants while preserving essential ionic composition. The successful adaptation enabled production scaling without dependence on commercial seawater sources.

#### Nutrient Supplementation Optimization

Enhanced nutrient supplementation was implemented to optimize phage production yields. Standard Pro99 nitrogen and phosphorus concentrations were increased to 2× normal levels. This enhancement was based on evidence that cyanophage infection substantially increases nutrient demands, with nitrogen uptake increasing immediately after infection and phosphorus assimilation rates increasing up to 8-fold within minutes of infection (Stent and Maaløe, 1953; Waldbauer et al., 2019). Nutrient limitation can severely compromise phage productivity, with phosphorus depletion causing delayed latent periods and up to 80% reduction in burst size (Wilson et al., 1996; Howard-Varona et al., 2024), while severe nutrient stress can lead to pseudolysogenic responses where infected cells fail to lyse (Rihtman, 2016; Cheng et al., 2019). Additionally, nitrogen limitation can completely prevent successful phage adsorption and replication in some cyanophage-host systems (McKindles, 2017). The 2× supplementation provided more stable growth patterns, higher maximum cell densities, reduced frequency of culture crashes, and support for the intensive biosynthetic demands of phage replication compared to standard nutrient levels.

### 2.2. Scaled Cultivation Systems

#### Vessel Preparation and Aeration System Design

All cultivation vessels were thoroughly cleaned and sterilized to prevent contamination during extended culture periods. Due to the high sensitivity of Prochlorococcus strains to trace metal concentrations (Moore et al., 2007; Chisholm Laboratory, 2022), special care was taken in preparing culture vessels. Vessels were acid-washed by soaking for 5 days in milliQ water containing trace metal grade hydrochloric acid (37 wt.% in H O, 99.999% trace metals basis, #339253-500ML, Sigma-Aldrich), followed by five rinses with milliQ water to remove acid residues. Intermediate-scale cultivation (1-8 L) was developed by adapting recommendations from (Chisholm Laboratory, 2022). We used PYREX® Round Media Storage Bottles (1395-1L and #1395-2L, Corning Incorporated) that were autoclaved three times with milliQ water at 121°C for 45 minutes to eliminate acid residues **(Supplementary Figure S1A)**. Large-scale operations (10-40 L) employed 12 L food-grade polyethylene terephthalate (PET) carboys that were sterilized by rinsing with 70% ethanol after acid washing and water rinsing, then allowed to dry in a sterile environment **(Supplementary Figure S2A)**.

Scaled cultivation required gentle aeration systems to provide CO supply and mixing while avoiding shear stress that could damage *Prochlorococcus* cells (Moore et al., 2007; Chisholm Laboratory, 2022). Aquarium pumps (Tetra® Whisper series) delivered filtered air at 0.1-0.2 L/min through silicone tubing fitted with Acro™ 37 TF vent devices (#4464, Cytiva) for sterilization. Custom cap modifications accommodated air inlet, outlet, and sampling ports: intermediate-scale bottles used Corning® PBT caps (#1395-45DC) with three holes for PTFE tubing **(Supplementary Figure S1B)**, while large-scale carboys utilized modified Jaece Industries Identi-plug™ Plastic Foam Stoppers (#L800-D) to allow tubing passage while maintaining airtight seals **(Supplementary Figure S2B)**. All tubing and cap components except foam stoppers were acid-washed, rinsed, and sterilized by dry-cycle autoclaving at 121°C for 30 minutes before assembly.

### 2.3. Phage Infection and Propagation

#### Host Culture Preparation and Timing

T7-like lytic *Prochlorococcus* phage P-SSP7 (Moran et al., 2005) was obtained from The Chisholm Lab at Massachusetts Institute of Technology. Successful phage infection required precise timing of host culture harvest at mid-exponential growth phase, corresponding to *P. marinus* MED4 cell densities of (0.1-1)×10 cells/mL as determined by flow cytometry or calibrated optical density measurements. This growth stage provided optimal host cell viability and metabolic activity for efficient phage replication.

#### Infection Protocol Optimization

Multiplicity of infection (MOI) was optimized based on experimental objectives. For routine maintenance and propagation, low MOI values (<0.01) were used following established protocols for cyanophage cultivation (Wilson et al., 1996; Zborowsky and Lindell, 2019; Laurenceau et al., 2021). This approach ensures gradual culture lysis, prevents premature host population depletion, and maximizes phage yield by allowing multiple rounds of infection before complete host exhaustion (Jo et al., 2024). For scaled production, MOI values of 0.5-1 × 10 ² provided optimal balance between infection efficiency and yield (Jo et al., 2024; Kosznik-Kwaśnicka et al., 2024; Wiebe et al., 2024).

Phage inoculum was prepared using standard lysate preparation methods (Wilson et al., 1996; Mruwat et al., 2021): fresh lysate was cleared of cellular debris by centrifugation at 8,000 × g for 10-15 minutes at 4°C, followed by 0.2 μm filtration to remove intact cells while preserving phage particles. Following phage addition, aeration was suspended for 60 minutes to facilitate phage adsorption without mechanical disruption.

Cultures were maintained under continuous light to maximize host metabolic activity and phage production. Culture lysis was monitored by visual inspection and optical density measurements, with progression from healthy green cultures to chlorotic, lysed cultures indicating successful infection (**Supplementary Figures S1C,D**). Optional nutrient supplementation (1× Pro99 nitrogen and phosphorus) every 3-4 days supported host metabolism during infection.

Complete lysis typically occurred within 6-9 days post-infection.

### 2.4. Multi-Stage Purification Protocols

#### Lysate Clearing by Removal of Host-Cell Debris

Following complete host cell lysis, crude phage lysates required clarification to remove cellular debris and unlysed cells, typically accomplished through centrifugation and/or microfiltration (Wilson et al., 1996; Mruwat et al., 2021). For small-scale routine phage propagation, lysates were cleared by centrifuging in 50 mL tubes at 8,000 × g for 15 minutes at 4°C followed by filtration of the supernatant through 0.22 μm PES membrane filters. The filtrate can be stored at 4°C, assessed for phage counting, and used to propagate or scale-up infection.

For large-scale infections, lysates were centrifuged at 26,000 × g using a J-LITE® JLA12.500 rotor (#C55767, Beckman Coulter) at 4°C for 30-40 minutes to pellet cellular debris while retaining phage particles in the supernatant. Cleared supernatants were filtered using Corning® 1000 mL Vacuum Filter/Storage Bottle Systems with 0.22 μm pore size, 54.5 cm² PES membrane (#431098, Corning Incorporated, USA), with multiple filtration units operated in parallel. Filtered lysates were pooled in sterile storage vessels. During optimization of the large-scale production pipeline, two additional treatments were introduced into the debris removal protocol. Lysates were pre-treated with 0.2 U/mL DNase I (30 minutes at room temperature) to reduce viscosity, facilitate clearing, and minimize phage losses (Yang et al., 2022), followed by NaCl addition to 2 M final concentration (#71376-5KG, Sigma-Aldrich, Inc., USA) with gentle stirring an 60-minute incubation at 4°C to facilitate phage particle release from membrane vesicles and cellular debris.

#### Phage Concentration Using Tangential Flow Filtration (TFF)

For intermediate-scale processing (1-8 L lysates), tangential flow filtration using a molecular weight cut-off (MWCO) 100 kDa membrane was used for initial phage concentration as 100 kDa provides a nice balance between retaining phage particles while allowing smaller proteins to pass through, enabling effective cross-flow velocity and transmembrane pressure to achieve a volumetric concentration factor of 8-10 (Grzenia et al., 2008; Malik et al., 2023). Vivaflow® 200 Tangential Flow Filtration Cassettes with Hydrosart® (HY) MWCO 100 kDa membrane (#VF20H4, Sartorius AG, USA) were assembled according to manufacturer specifications, with careful attention to flow rates and pressure limits to prevent membrane fouling and phage damage. Between processing batches, TFF cassettes were cleaned by recirculating 0.5 M NaOH for 30 minutes, followed by thorough washing with autoclaved deionized water and sterilization with 70% ethanol before final rinsing.

#### Phage Concentration Through Polyethylene Glycol (PEG) Precipitation and Sucrose Cushion Ultracentrifugation

PEG precipitation in the presence of NaCl provides effective phage concentration applicable across all scales (Yamamoto et al., 1970). For each liter of cleared lysate, 116 g NaCl (#71376-5KG, Sigma-Aldrich, Inc., USA) and 100 g PEG 8,000 (#89510-1KG-F, Sigma-Aldrich) were added sequentially with gentle stirring at 4°C, followed by overnight incubation. PEG-bound phages were recovered by centrifugation at 26,000 × g for 25 minutes at 4°C and the supernatant was carefully decanted. The optimized protocol included careful removal of remaining PEG solution by inverting bottles in a sterile cabinet for 5-10 minutes and wiping walls with sterile tissue. The dry pellets were resuspended in 10 mL cold SM buffer (100 mM NaCl, 8 mM MgSO, 50 mM Tris-HCl, pH 7.5) through overnight orbital shaking (∼100 rpm) at 4°C, followed by gentle pipetting to complete resuspension.

Sucrose cushion ultracentrifugation provided efficient further concentration and partial purification (Mbiguino and Menezes, 1991), with the advantage of processing larger phage solution volumes (up to 50 mL per tube) compared to subsequent CsCl gradients. A 38% sucrose solution (#S7903-5KG, Sigma-Aldrich, Inc., USA) was prepared in SM buffer and filter-sterilized through 0.22 μm filters (Hurwitz et al., 2012). The resuspended phage solution (40-50 mL) was added to 70 mL Polycarbonate Bottles (#355622, Beckman Coulter, Inc., USA), then 20-30 mL of 38% sucrose solution was carefully underlayered using sterile 9-inch Pasteur pipets (#7095D-9, Corning Incorporated, USA) to form a distinct density interface. Tubes were centrifuged using Ultracentrifuge Rotor Type 45 Ti (#339160, Beckman Coulter, Inc., USA) at 113,000 × g for 3.5 hours at 4°C. Following centrifugation, the supernatant was carefully decanted without disturbing the phage pellet. Remaining solution droplets were carefully removed using sterile KimWipe paper. Pellets were air-dried for 10-15 minutes in a sterile cabinet, then resuspended in 2 mL SM buffer through overnight orbital shaking at 80-100 rpm at 4°C followed by gentle pipetting with wide-orifice tips to minimize shear forces.

#### Cesium Chloride (CsCl) Density Gradient Purification: Single-Step and Two-Step Protocols

Final purification utilized cesium chloride density gradient centrifugation to separate phage particles from remaining contaminants based on particle density (Bachrach and Friedmann, 1971). CsCl solutions (#562599, Sigma-Aldrich, Inc., USA) were prepared in SM buffer at specific densities and filter-sterilized through 0.2 μm filters. The protocol, modified from (Hurwitz et al., 2012), used step gradients with solutions at densities of 1.65, 1.55, 1.5, 1.4, and 1.2 g/mL prepared by dissolving appropriate amounts of CsCl in SM buffer. Gradients were constructed in 14 mL Open-Top Thinwall Ultra-Clear Tubes (#344060, Beckman Coulter), layering from highest to lowest density, with approximately 3 mL of concentrated phage solution carefully overlaid. Following centrifugation in a SW 40 Ti Swinging-Bucket Rotor (#330070, Beckman Coulter) at 154,000 × g for 3.5 hours at 4°C, phage bands visible at the 1.4-1.5 g/mL interface were collected using a 3 mL syringe with 20-gauge needle by careful side puncture.

For enhanced purity, a second gradient step was performed by mixing collected phage-containing CsCl solution with fresh 1.5 g/mL CsCl solution to achieve 10 mL total volume, overlaying this onto 1 mL of 1.65 g/mL CsCl solution, and centrifuging at 98,000 × g for 12-16 hours at 4°C (Bachrach and Friedmann, 1971).

#### CsCl Removal by Stepwise Dialysis

Stepwise dialysis was used to remove CsCl from purified phage preparations while reducing osmotic pressure and avoiding activity loss (Luong et al., 2020; Carroll-Portillo et al., 2021). The process employed 3 mL Slide-A-Lyzer™ G3 Dialysis Cassettes with 10K MWCO membrane (#A52981, Thermo Fisher Scientific). Each dialysis step was conducted for 30-60 minutes at 4°C in 1 L beakers under gentle magnetic stirring. The dialysis sequence proceeded as follows: initial dialysis against 300 mL modified SM buffer containing 5 M NaCl, followed by stepwise dilution with fresh SM buffer (adding 200 mL to reduce to ∼3 M NaCl, then 500 mL to reduce to ∼1.5 M NaCl). Subsequently, 800 mL spent buffer was removed and replaced with 200 mL fresh buffer (reducing to ∼0.5 M NaCl), followed by transfer to 1 L fresh SM buffer, and finally transfer to fresh 1 L SM buffer for overnight dialysis at 4°C.

### 2.5. Host Cell and Phage Quantification Methods for Accurate and Rapid MOI Control

#### Flow Cytometry-Based Cell Enumeration and Optical Density Calibration

Flow cytometry provides precise absolute cell enumeration of host cells (Chisholm et al., 1988). A BD Influx Fluorescence Activated Cell Sorter (FACS, BD Biosciences, San Jose, CA) equipped with a 100 μm nozzle was used. Prior to analysis, the instrument was optimized and calibrated using 3 μm Ultra Rainbow Fluorescent Particles (Spherotech, Lake Forest, IL) to ensure consistent performance.

Given the small size (∼0.6 μm diameter) and fragile nature of *P. marinus* MED4 cells, a sequential gating strategy was employed to distinguish viable cells from debris and non-viable particles. Cells were first stained with SYBR Gold nucleic acid stain and analyzed using 488 nm excitation and 530/30 nm emission to identify DNA-containing particles. DNA-positive cells were then analyzed for chlorophyll autofluorescence using 640 nm laser excitation and 670/30 nm emission to specifically identify photosynthetically active *P. marinus* cells.

To enable rapid cell quantification during scaled operations, a calibration curve correlating optical density at 750 nm with absolute cell counts was established using 25 independent *P. marinus* MED4 culture samples with OD measurements ranging from 0.03 to 0.26. Parallel OD measurements (Genesys 20 Spectrophotometer, Thermo Fisher Scientific) and flow cytometry-based absolute counts were performed, with data fitted to a linear regression model with 95% confidence intervals (R² = 0.935, p < 0.001). The resulting calibration equation (*P. marinus* = 3.4 × 10 × OD + 9.5 × 10) enabled precise multiplicity of infection calculations essential for reproducible phage infections across production scales.

#### Fluorescence Microscopy-Based Phage Enumeration

Total phage particle quantification utilized fluorescence microscopy following nucleic acid staining to visualize individual virions, conducted with modifications to the method reported earlier (Patel et al., 2007). To ensure counting of only intact phage particles with protected DNA, samples were pre-treated with DNase I to degrade exogenous DNA. For each sample, 100 μL of phage suspension was treated with 10 μL reaction buffer and 10 μL DNase I (1 U/μL) from the Thermo Scientific DNase I kit (#EN0521) at room temperature for 60 minutes, followed by inactivation with 10 μL EDTA stop solution.

Following DNase treatment, phage particles were stained with 2 μL of 100× SYBR Gold nucleic acid stain (#S11494, Invitrogen) in the dark for 20 minutes. Samples were diluted to 1 mL with 0.1 μm filtered PBS and filtered through 25 mm, 0.02 μm pore size Cytiva Whatman™ Anodisc™ filter discs (#09-926-34) using vacuum filtration. Filter discs were washed three times with filtered PBS, air-dried, and mounted on microscope slides with 50% glycerol. Slides were examined using a Leica epifluorescence microscope with 100× oil immersion objective and appropriate filters for SYBR Gold visualization (488 nm excitation, 525 nm emission). Approximately 20 random fields were captured per sample for statistical reliability, with image analysis performed using Fiji (ImageJ) software for automated particle counting.

#### Most Probable Number (MPN) Assay for Infectious Titer

The MPN assay quantified infectious phage particles through statistical analysis of dilution series infections, performed using serial dilutions based on (Suttle and Chan, 1993; Faulhaber et al., 2012; Chénard and Chan, 2018). This approach provided advantages over traditional plaque assays for cyanophages, including higher throughput and better detection of infectious particles that may not form clear plaques.

Serial dilutions of phage samples were prepared in 96-well plates using Pro99 medium with 2× nitrogen and phosphorus concentrations. Initial dilutions began at 10 ³ concentration, with ten-fold serial dilutions extending to 10 ¹ . Using multi-channel pipettors, 30 μL of exponentially growing MED4 host cells was added to test wells, followed by 20 μL of each dilution level, with final volume adjusted to 250 μL with additional Pro99 medium. Following 1-hour phage adsorption at appropriate light and temperature conditions, plates were sealed and incubated for up to 2 weeks with regular monitoring. Wells showing significant reduction in chlorophyll fluorescence compared to controls were scored as positive for lysis using a Biotek Cytation C10 plate reader with 485 nm excitation and 675 nm emission filters. MPN calculations were performed using standard statistical tables or software to determine infectious units per mL in original samples. MPN assays were performed using 6 replicates for every dilution, with statistics calculated according to (Jarvis et al., 2010). Calculation of MPN estimates and statistics was performed using the MPN R package (Ferguson and Ihrie, 2024).

### 2.6. Validation of Phage Purity and Particle Density through Cryo-EM

Visual verification using cryo-electron microscopy was used as a final validation step for the virus purification protocol at each stage of the scale-up and for each round of optimization. Briefly, 3 µL of the final virus suspension was loaded onto glow-discharged holey carbon grids (Quantifoil Q2/2 or 2/1, 200 or 300 mesh) and blotted for 2.5–3.5 s. The samples were frozen in a vitreous layer of ice by rapidly plunging in liquid ethane on a Leica EMGP2, followed by storage in liquid nitrogen. For screening and imaging, grids were loaded on a 300 keV TFS Titan Krios G3i cryo-electron microscope. Grids were imaged on a Gatan K3 (Gatan Inc.) camera using a Bioquantum energy filter, with the slit width at 20 eV, at 81,000× nominal magnification and calibrated pixel size of 1.1 Å, with a total dose of ∼45 e^-^/Å^2^/s at a nominal defocus of -0.75 to - 1.75 µm. Dose optimization was not performed for images collected at lower magnifications. Large scale datasets were collected using standard software like TFS EPU or SerialEM and image processing was carried out using CryoSPARC v 4.7.1 (Mastronarde, 2005). Reference free 2D classification was used to sort the virus particles into uniform classes. Details of image processing as well as subsequent 3D refinement and high-resolution structure of P-SSP7 will be part of a follow-up manuscript.

### 2.7. Validation of Phage Infectivity through Cryo-ET of Host-Phage Interactions

**(A)** *P. marinus* MED4 cells (∼10^9^ cells/mL) were infected with phage P-SSP7 at an MOI of 40 to synchronize infection of all cells for visualizing host-phage interaction using cryo-electron tomography (cryo-ET). A high MOI has been used in cryo-ET studies of host-phage interactions (Murata et al., 2017). At intervals of 15 mins and 45 mins, 3 µL aliquots from the bulk infection were mixed with 10 nm BSA coated colloidal gold fiducials (Aurion SKU: 200.133) and were loaded onto holey carbon grids (Quantifoil Q2/2 or 2/1, 200 or 300 mesh). Grids were rested horizontally on a benchtop in tweezers for 2–3 mins under ambient conditions and excess suspension was blotted for 1.5–2.5 s. Grids were vitrified by plunge freezing in liquid ethane using a Leica EM GP2. Tomographic data were collected on the TFS Titan Krios G3i on a Gatan K3 (Gatan Inc.) camera using a Bioquantum energy filter, with the slit width at 20 e^-^V, using a Volta Phase Plate, at 33,000 x nominal magnification. Tilt series were collected from -/+54° at 3° intervals using SerialEM. Tilt images were recorded as movies with 10-12 subframes and an exposure of 1.02 s in super resolution mode at a pixel size of 1.3 Å and a total dose of 110-120 e^-^/Å^2^ and a nominal defocus of -4 µm. Custom in-house scripts were used to run the automated workflow of image processing from motion correction to final 3D tomogram calculation. MotionCorr2 (Zheng et al., 2017) was used to perform motion correction of the individual movies. Restacking and CTF correction and reconstruction of the tomograms were performed using IMOD (Kremer et al., 1996) and ETomo suite (Mastronarde and Held, 2017). Final reconstructed tomograms were binned 8x relative to the raw data and visualization of the 3D volumes was done using 3dmod.

## 3. Results

### Systematic Scale-Up Strategy and Progressive Optimization

Our scale-up approach systematically progressed from laboratory maintenance volumes (5-100 mL) through intermediate development scales (1-8 L) to production-scale quantities (10-40 L), with each phase providing critical insights that informed subsequent optimization. This iterative development strategy, guided by quality assessment at each scale, ultimately achieved the major improvements in sample quality necessary for routine structural studies.

### 3.1. Small-Scale Foundation (5-100 mL): Establishing Baseline Protocols

The small-scale system established fundamental protocols for MED4 cultivation and P-SSP7 propagation using standard laboratory equipment (**Figure 1**). Routine maintenance was successfully performed using Falcon® 5 mL test tubes for volumes up to 10 mL and CELLSTAR® filter cap flasks for 25-100 mL cultures (**Figure 1A**). This scale enabled consistent culture transfers every 2-3 weeks during mid-exponential growth phase, establishing reliable growth patterns and infection kinetics essential for larger-scale operations.

**Figure 1.**
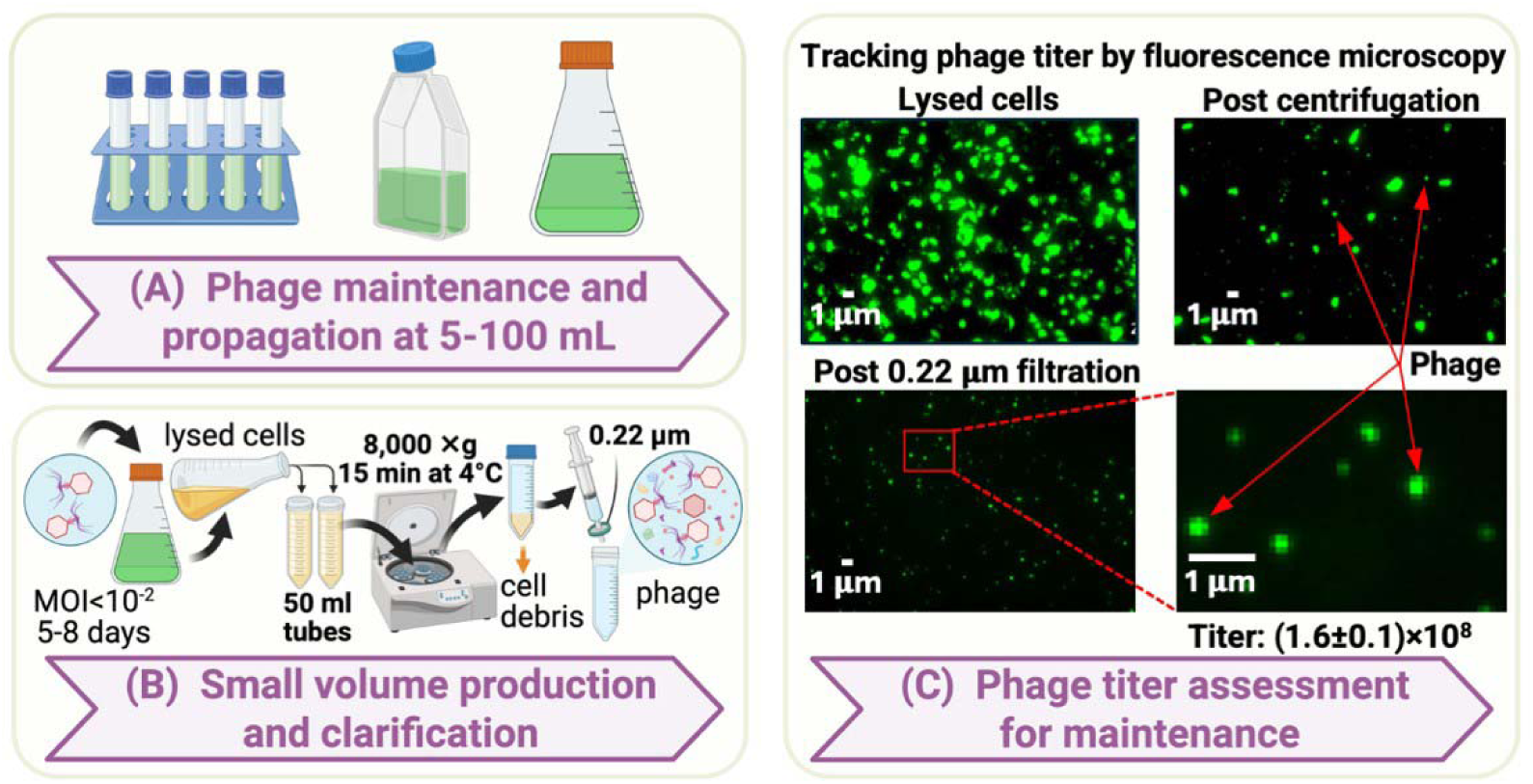
Small-scale *Prochlorococcus marinus MED4* cultivation and cyanophage P-SSP7 propagation workflow. **(A)** Host culture maintenance systems for 5-100 mL volumes using test tubes and filter cap flasks. **(B)** Infection monitoring and lysis progression showing culture clearing and phage harvest procedures. **(C)** Fluorescence microscopy-based phage quantification using SYBR Gold staining and automated particle enumeration workflow. *Created with BioRender.com*

Phage infection protocols were optimized using MOI values below 0.01 leading to gradual culture lysis over 6-9 days (**Figure 1B**). The infection process consistently resulted in complete culture clearing, confirmed by visual inspection and flow cytometry analysis. Simple lysate harvest involved centrifugation at 8,000 × g for 15 minutes followed by 0.2 μm filtration yielded total phage titers of (1.6±0.1)×10 particles/mL as determined by fluorescence microscopy (**Figure 1C**). This total count includes both infectious and non-infectious particles and should be interpreted cautiously for estimating harvestable phage yields. The ratio of infectious to total particles can vary dramatically—from nearly 100% to less than 1%—depending on cultivation and environmental conditions (Bratbak et al., 1998; Bhat et al., 2022). For P-SSP7 specifically, elevated light intensity increases mispackaged particles from 4% under low light to nearly 90% under high light conditions, though co-cultivation with bacterial partners like *Alteromonas* can reduce mispackaging to 3% even at high light (Laurenceau et al., 2021). Additionally, purification steps progressively reduce infectious titers through particle inactivation. Studies with *E. coli* phages demonstrate that PEG precipitation alone reduces activity by 5-50% depending on PEG molecular weight average, while CsCl gradient purification can decrease infectivity by 40-95% (Carroll-Portillo et al., 2021). Consequently, infectious titers were not quantified at this foundational scale, with more rigorous infectivity assessments implemented at production scales where purification optimization became critical.

An important methodological advance at this scale was the development of the optical density-based enumeration method for host cells (**Figure 2**). Given the small size (∼0.6 μm diameter) and fragile nature of *P. marinus* cells, this approach required establishing a rigorous flow cytometric gating strategy to distinguish viable, photosynthetically competent cells from debris and non-viable particles. Sequential gating first identified DNA-containing cells using SYBR Gold staining (**Figure 2A**), then specifically targeted photosynthetically active *P. marinus* cells based on their characteristic chlorophyll autofluorescence signature (**Figure 2B**). The calibration curve established using 25 independent MED4 culture samples provided reliable cell density estimates from OD measurements (**Figure 2C**), enabling precise MOI calculations essential for reproducible infections for scaled phage production and structural studies. This rapid quantification method proved indispensable for efficient workflow management during larger-scale operations.

**Figure 2.**
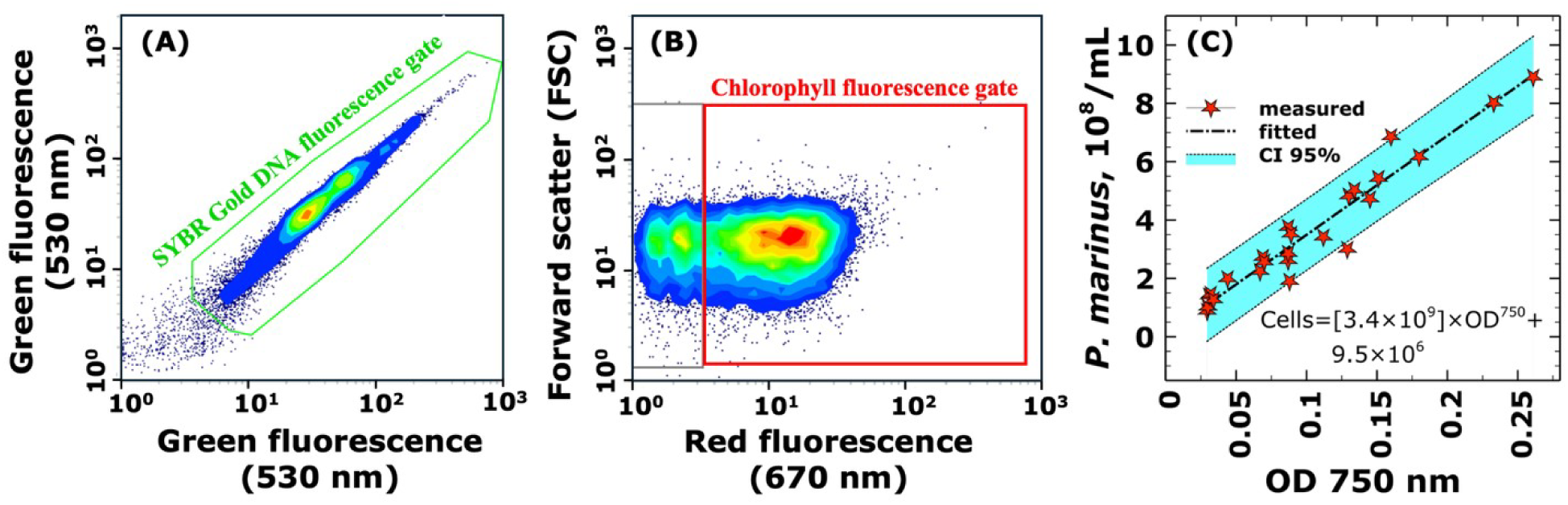
Flow cytometric identification of viable *Prochlorococcus marinus* MED4 cells and optical density calibration for accurate cell enumeration. **(A)** Dual-parameter analysis of SYBR Gold-stained cells (488 nm excitation, 530/30 nm emission) identifying DNA-containing cells (green gate) and excluding cellular debris and particles lacking intact genetic material. **(B)** Forward scatter versus chlorophyll autofluorescence (640 nm excitation, 670/30 nm emission) analysis of DNA-positive cells from panel A, specifically targeting photosynthetically active *P. marinus* MED4 cells (red gate) based on their characteristic chlorophyll fluorescence signature. **(C)** Linear correlation between optical density at 750 nm and absolute counts of viable, photosynthetically competent *P. marinus* cells determined by the sequential gating strategy for 25 independent culture samples (R² = 0.935, p < 0.001). Red symbols represent individual measurements with linear regression fit (dashed line) and 95% confidence intervals (shaded region). The calibration equation *P. marinus* = 3.4 × 10 × OD + 9.5 × 10 enables rapid enumeration of viable cells for precise multiplicity of infection calculations in phage infection studies, ensuring accurate assessment of host cell availability for viral replication.

### 3.2. Intermediate-Scale Development (1-8 L): Identifying Key Challenges

The intermediate scale successfully demonstrated local seawater adaptation and custom cultivation systems through systematic optimization (**Figure 3, Supplementary Figure S1C,E**). Systematic adaptation from commercial Sargasso Seawater to local Salish Sea seawater was completed over two months without significant loss of culture viability. Enhanced nutrient supplementation (2× nitrogen and phosphorus) provided stable growth patterns and higher *P. marinus* cell densities of 0.5-1 × 10 cells/mL in 1-2 L PYREX bottles (**Figure 3A, Supplementary Figure S1C**) suitable for efficient phage infection, providing abundant infection targets for maximum phage yield.

**Figure 3.**
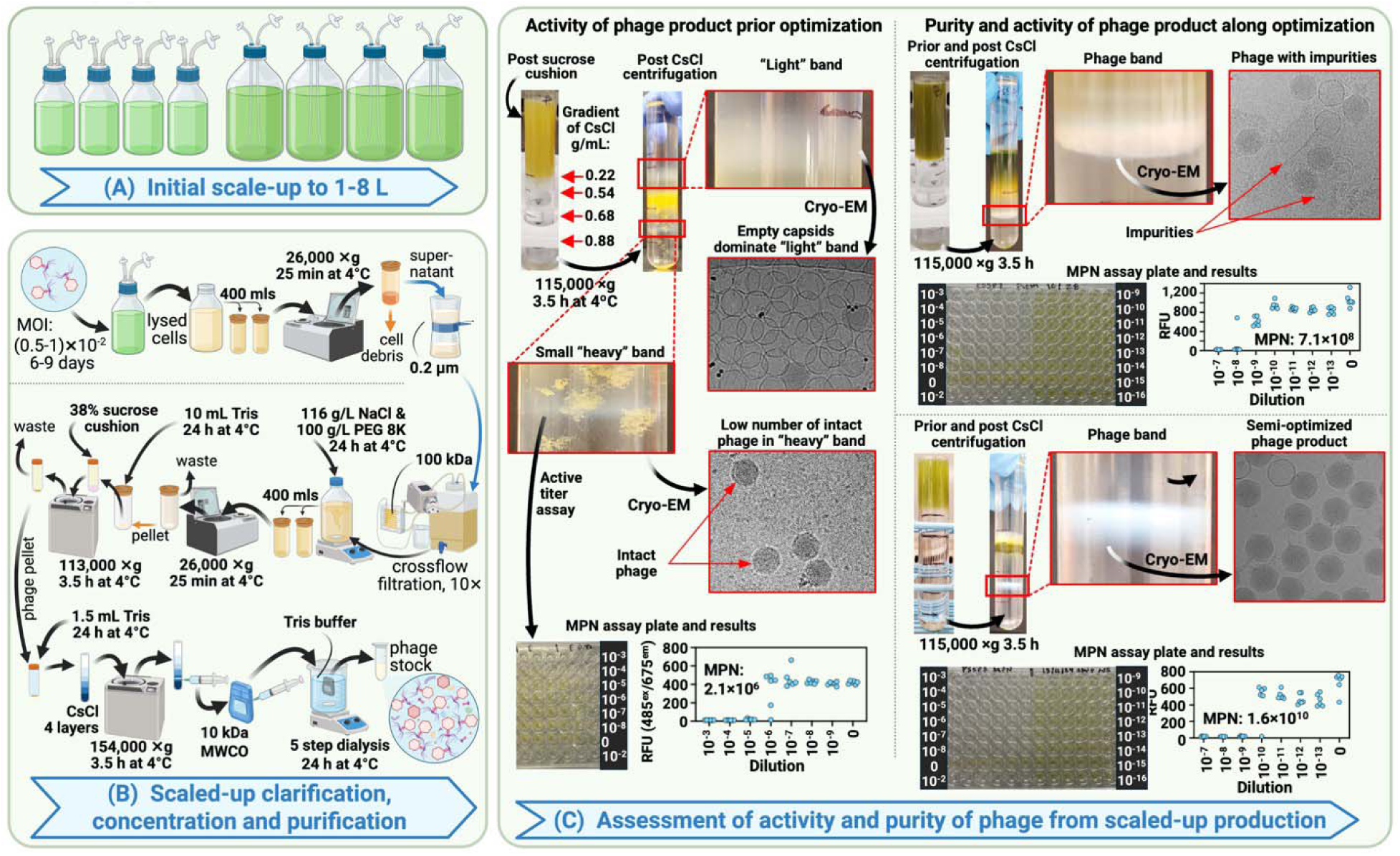
Intermediate-scale development revealing cultivation success and purification challenges (1-8 L). **(A)** Custom cultivation system with PYREX bottles and modified caps providing sterile aeration and sampling access. **(B)** Complete purification workflow including tangential flow filtration, PEG precipitation, and ultracentrifugation with quantitative yield data. **(C)** Quality assessment by MPN assay and cryo-EM validation revealing significant debris contamination requiring optimization. *Created with BioRender.com*

However, intermediate-scale development required iterative optimization through three distinct phases (**Figure 3B,C**). Initial scaling from cell flasks to 1-1.5 L culture volumes in 1 L PYREX bottles without aeration yielded MPN titers of only 2.1×10 infectious units/mL—insufficient for structural applications and limited by low culture volumes and suboptimal cultivation conditions. The second phase involved scaling to 4-8 L total culture volumes using multiple parallel 2 L vessels with custom aeration systems (**Figure 3A, Supplementary Figure S1E**). Gentle aeration at 0.1-0.2 L/min maintained adequate CO supply and mixing without cell damage, while optimized host cultivation densities increased infectious titers to 7.1×10 units/mL. The final phase of intermediate optimization refined infection conditions through additional nutrient supplementation during infection, optimized infection timing before phage harvesting, and enhanced purification workflows. This comprehensive approach achieved MPN titers of 1.6×10¹ infectious units/mL.

Along with directing phage titer optimization, cryo-EM validation at each stage guided purification improvements and revealed progressive enhancement in sample quality (**Figure 3B,C**). Initial preparations from 1-1.5 L cultures suffered from extensive debris contamination and the problematic two-band CsCl separation, but systematic refinement of harvesting procedures, PEG precipitation parameters, and ultracentrifugation conditions progressively reduced contaminants while improving phage recovery (**Figure 3B**). The optimized purification workflow combining tangential flow filtration, enhanced PEG precipitation, sucrose cushion ultracentrifugation, and CsCl density gradient centrifugation ultimately yielded total recoverable infectious units of 5×10¹ .

However, despite these notable improvements in both titer and purity, cryo-EM validation revealed persistent limitations for high-resolution structural applications (**Figure 3C**). Sample preparations exhibited only 6-8 virus particles per field of view at 11,500× magnification, accompanied by residual debris contamination insufficient for effective collection of the extensive imaging datasets required for comprehensive structural determination. This particle density limitation becomes particularly critical when resolving symmetry-mismatched features like the portal-tail complex, which require significantly larger datasets due to the inability to apply symmetry averaging during reconstruction. This cryo-EM-guided intermediate-scale experience demonstrated that achieving structural study quality would require not only further volume scaling but also implementing fundamentally enhanced purification strategies specifically designed to meet the stringent particle density and purity requirements essential for high-resolution structural studies.

### 3.3. Large -Scale Optimization (10-40 L): Achieving high phage particle density for high resolution structural studies

Building on lessons learned from intermediate-scale work, the large-scale system incorporated several critical improvements that ultimately achieved the quality improvements necessary for routine structural studies (**Figure 4**). Twelve-liter PET carboys enabled cultivation of 10 L cultures per vessel, with multiple carboys operated in parallel to achieve 20-40 L total working volumes while maintaining the established seawater adaptation and enhanced nutrient supplementation protocols (**Figure 4A, Supplementary Figure S2A,C,D**). Further optimization of culture scaling, custom-designed aeration systems, nutrient supply timing, and infection parameters consistently achieved infectious phage titers of 3.1×10¹² units/mL, representing 100-to1000-fold improvements over intermediate-scale preparations (**Table 1, Figure 4C**).

**Figure 4.**
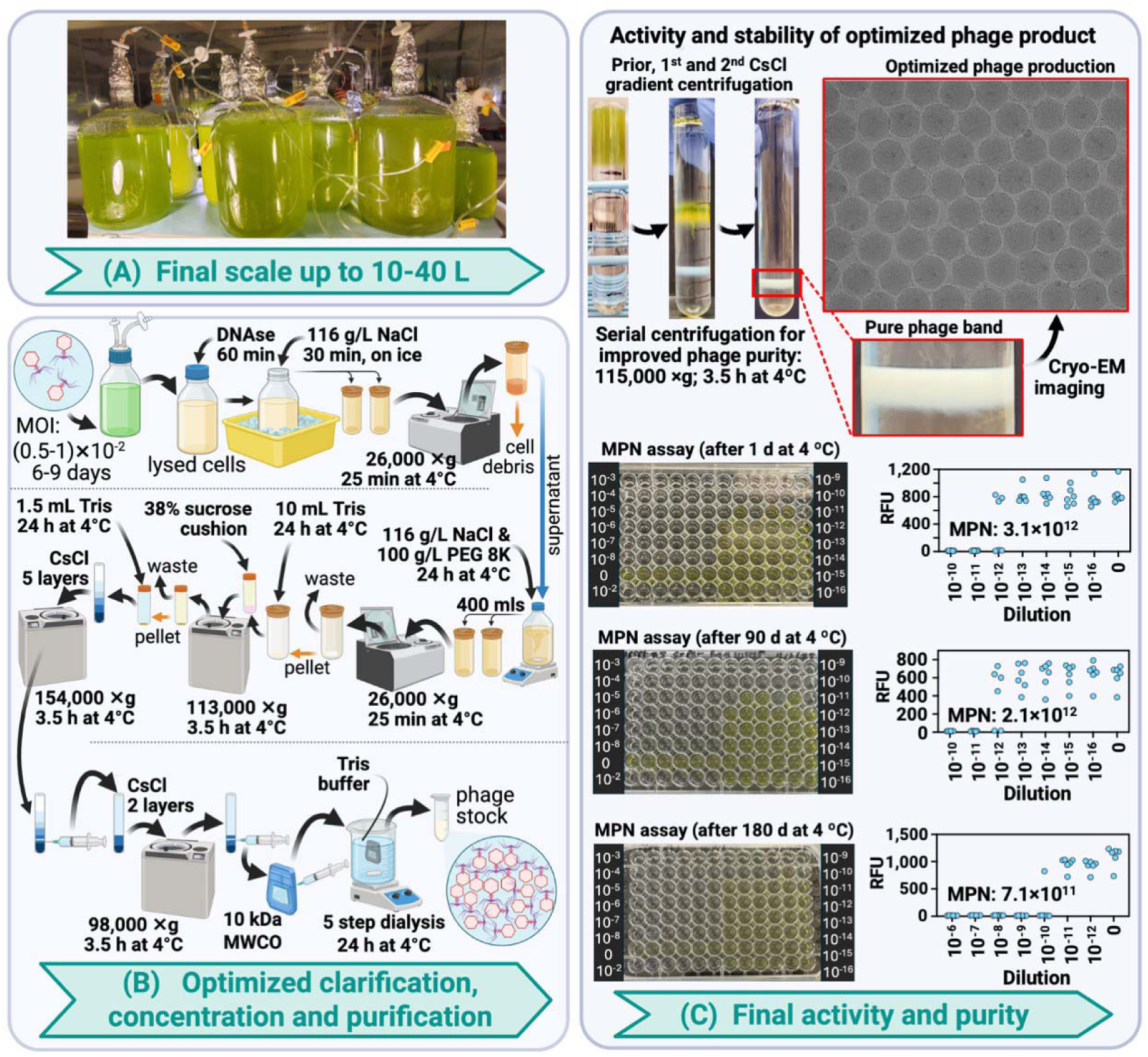
Large-scale phage production system achieving structural study quality (10-40 L). **(A)** Scaled cultivation using 12 L PET carboys with optimized aeration for parallel processing of maximum volumes. **(B)** Enhanced purification workflow featuring direct PEG precipitation and two-step CsCl density gradient centrifugation for superior purity. **(C)** Final preparation characterization showing improved phage yields, infectivity, and cryo-EM validation confirming structural study suitability. *Created with BioRender.com*

**Table 1.** Comparative production performance and applications across development scales.

| Scale | Culture volume (L) | Vessel type | Seawater source | Infectious titer ± CI<sup>1</sup> (MPN units/ml) | Total infectious units recovered ± CI (MPN units) | Application of the virus suspension |
| --- | --- | --- | --- | --- | --- | --- |
| Small | 0.05-0.1 | Cell Culture Tubes or Flasks | Commercial Sargasso Seawater | NA | $1.6 \pm 0.1 \times 10^4$ | Routine maintenance, initial propagation |
| Intermediate initial | 1-1.5 | 1 L PYREX bottles | Local adapted Salish Seawater | $(2.1 \pm 2.3) \times 10^6 - (7.1 \pm 4.2) \times 10^8$ | $(2.9 \pm 3.2) \times 10^6 - (9.9 \pm 5.9) \times 10^8$ | Phage production method development, optimization |
| Intermediate optimized | 4-8 | 2 L PYREX bottles with custom aeration | Local adapted Salish Seawater | $(1.6 \pm 1.7) \times 10^{10}$ | $(5 \pm 5.3) \times 10^{11}$ | Purification method development, optimization |
| Large optimized, freshly harvested | up to 40 | 12 L PET Carboys | Local adapted Salish Seawater | $(3.1 \pm 3.1) \times 10^{12}$ | $(1 \pm 1) \times 10^{13}$ | High-resolution structural studies |
| same, stored 3 months at 4°C | same | same | same | $(2.1 \pm 2.3) \times 10^{12}$ | $(6.8 \pm 7.4) \times 10^{12}$ | same |
| same, stored 6 months at 4°C | same | same | same | $(7.1 \pm 4.2) \times 10^{11}$ | $(2.3 \pm 1.4) \times 10^{12}$ | same |

<sup>1</sup> Confidence Intervals (CI) reflect the inherent statistical uncertainty of MPN methodology, which estimates viable counts through 539 dilution series. Wide intervals (typically $\pm 1-2$ log units) are characteristic of this statistical approach, not experimental error (Jarvis et al., 2010).

The most significant advances came through systematic optimization of phage harvesting and purification workflows specifically designed to meet cryo-EM quality requirements (**Figure 4B**). Key modifications included elimination of tangential flow filtration in the final pipeline, implementing direct PEG precipitation instead, and most critically, implementation of two-step CsCl gradient purification for enhanced purity. The first CsCl gradient separated infectious particles from debris and empty capsids, while the second gradient step further purified the infectious phage band, substantially reducing contaminating material. Optimized stepwise dialysis protocols ensured complete CsCl removal while preserving viral integrity. This enhanced pipeline yielded total recoverable infectious units of 10¹³, with MPN assays confirming preserved structural integrity (**Figure 4C**).

Importantly, the final preparations demonstrated robust stability over extended storage periods. Freshly produced phage with initial MPN titers of 3.1×10¹² units/mL retained 68% activity (2.1×10¹² units/mL) after 3 months at 4°C, and 23% activity (7.1×10¹¹ units/mL) after 6 months. This relatively slow activity decline ensures consistent and reproducible imaging campaigns while providing sufficient material for follow-up experiments including infection dynamics studies and integrated multi-omics analyses.

#### Production Performance Summary

The complete scale-up achieved substantial increases in production volume (up to 800-fold from phage maintenance volumes of ∼50 mL to final 40 L scales) while substantially improving sample quality (**Tables 1**). Small-scale foundations (5-100 mL) using commercial seawater established baseline protocols with total phage titers in lysate of ∼10^8^ particles/mL suitable for routine maintenance. Intermediate-scale development (1-8 L) successfully demonstrated local seawater adaptation and cost reduction but revealed infectious titer and purification challenges limiting structural applications. Large-scale optimization (10-40 L) achieved final active phage titers exceeding 3×10^12^ infectious units/mL with >95% purity suitable for high-resolution structural studies, while significantly reducing production costs through local seawater adaptation.

### 3.4. Visual validation of the virus suspension using Cryo-EM Imaging

The progressive optimization achieved major improvements in sample quality as validated by the earlier cryo-EM analysis at each stage. **Figure 5** shows the four main preparation stages described above in a layout providing direct comparison. For the Intermediate Initial condition (**Figure 5A**), the overview image shows 8 P-SSP7 particles with intact DNA in the image along with extensive debris and several DNA-empty P-SSP7 particles. However, 4 of the 8 DNA-intact particles are clustered together and outside the typical fields of view used for high magnification data collection indicated by the black, red, and blue rectangles. Thus, a total of 4 intact particles are seen across the 3 high magnification areas that were imaged, yielding an average of about 1 per imaged field of view. A different arrangement of areas imaged per hole could have yielded anywhere between 0 to 8 particles per 3 imaged areas in this example for downstream processing. We noted that for dilute particle densities the P-SSP7 particles clustered near the edges of the carbon holes, so for consistency of comparison, we used the same relative areas for all dilute conditions. For more crowded conditions, we kept the same general layout but moved the center of the imaged areas closer to the center so each image largely avoided any carbon edge to maximize particle counts. The Intermediate Optimized sample (**Figure 5B**) is also dilute with most of the particles seen clustering near the TEM grid carbon edge. However, the average number of intact particles per high-magnification FOV are closer to 3. In contrast, the Large-Scale Initial (**Figure 5C**) and Large Scale Optimized (**Figure 5D**) samples show significantly more particles that span the hole rather than clustering near the edges. While the Large-Scale Initial (**Figure 5C**) high-magnification FOV images show an average of 39 intact particles, the Large-Scale Optimized (**Figure 5D**) sample has minimal debris contamination and shows particles adopting a close-packed monolayer which is near ideal for high-resolution cryo-EM data collection and results in an average of 66 intact particles per FOV. That improvement in particle density also means the Large-Scale Optimized sample would need to collect 66× fewer images to get the same final particle count as the original Intermediate Initial condition (**Table 2**). Importantly, each of the three high magnification images taken per hole all show similar particle numbers demonstrating improved efficiency for data collection.

**Figure 5.**
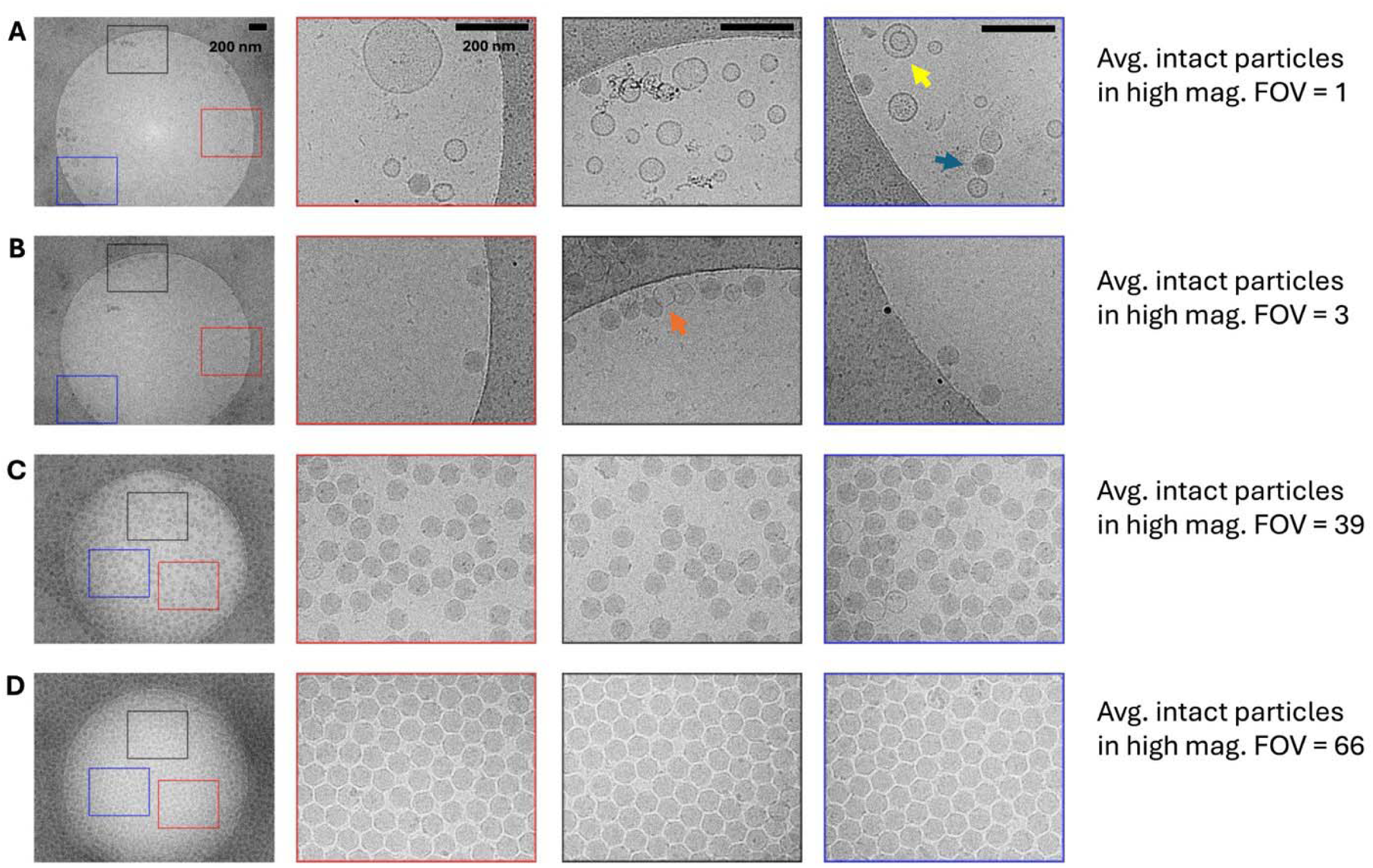
Cryo-EM validation demonstrating systematic quality improvement through iterative optimization. **(A,B,C,D)** Low magnification overview image (11,500×, scale bar 200 nm) tracking debris reduction across optimization stages with highlighted high-magnification field of view areas (red, blue, black borders) showing progressive improvement in particle density and quality. To the right of each overview image are the corresponding high-magnification FOV areas highlighted by the red, blue, and black boxes. **(A)** Initial preparation shows extensive cellular debris and numerous empty capsids. **(B)** Early optimization attempts with persistent contamination and low particle density. **(C)** Intermediate optimization showing reduced debris and increased intact particles. **(D)** Final optimized preparations displaying dense, ordered particle arrays suitable for high-resolution structural determination.

**Table 2.** Cryo-EM data collection efficiency and sample quality metrics across optimization stages.

| **Optimization stage** | **Average particles per image** | **Number of movies needed\*** | **Microscope time needed** | **Factor amount** | **Debris phages** | **Empty** |
| --- | --- | --- | --- | --- | --- | --- |
| Intermediate initial | 1 | 500,000 | 62.5 days | baseline | high | ~50% |
| Intermediate optimized | 3 | 166,667 | 20.8 days | 3x | reduced | ~20% |
| Large scale initial | 39 | 12,820 | 1.6 days | 39x | low | <5% |
| Large scale optimized | 66 | 7,575 | 1.0 day | >60x | minimal | <1% |

\* Typical single particle cryo-EM datasets usually need 100k-1000k particles for suitable 2D classification and 3D refinement so we used a mid-range of 500k for current target estimate.
\*\* The average readout rate for the K3 camera is ~8,000 images per day which is what we assume here. The table assumes perfect collection conditions, no instrument related downtime, no excluded images and all particles imaged are selected in the classification.

The most significant practical achievement of our optimization is the dramatic reduction in cryo-EM data collection time requirements (**Table 2**). Typical datasets for high-resolution single-particle reconstructions often require a few thousand to millions of equivalent “particles” at random Euler angles in the vitrified ice to generate structures with better than 4 Angstrom resolution. Larger particles, and particles with low symmetry or symmetry mismatch, generally require more particle copies. For our purpose, we chose an intermediate range of 500,000 particles as the target for comparison. For the Intermediate Initial stage of production with only 1 particle per average high magnification field of view, it would require 500,000 movies to be collected and would require 62.5 days of microscope access to complete the dataset (assuming full-time data collection at a rate of 8000 movies per day). This is clearly not feasible for most research groups. Our final optimized preparation, with 66 particles per FOV, would reduce the required microscope time from ∼62 days to ∼1 day for collecting 500,000 particles. Since most research groups only have limited access to cryo-EM facilities and usually allocate 24-72 hours per sample for data collection, our workflow makes comprehensive structural studies of marine cyanophages practically feasible for most research groups.

### 3.5. Visual validation of MED4 infection by P-SSP7 with cryo-ET

Building on the scalable production workflow that yielded high-titer, single-particle cryo-EM grade samples, we next evaluated whether these gains translate into reproducible, in situ visualization of infection by cryo-electron tomography (cryo-ET). Synchronized Prochlorococcus marinus MED4 cells were infected with P-SSP7 at an MOI of 40 and an aliquot was directly deposited onto a TEM grid shortly before plunge freezing. The resulting vitrified grids showed intact cells embedded in uniform vitreous ice and minimal extracellular debris. We then acquired several dose-symmetric tilt series on a 300 keV cryo-EM instrument and computationally reconstructed the 3D tomograms to capture host-phage interactions in 3D (**Figure 6**). Across independently prepared batches, tomograms consistently captured multiple stages of the adsorption process on single cells, including (i) free particles proximal to the cell surface, (ii) reversibly attached virions making initial contact, and (iii) tightly bound particles with the portal–tail axis oriented approximately normal to the membrane. The improved sample purity increased the fraction of tilt series suitable for downstream analysis. While a more rigorous time-resolved study is being performed separately to track the full range and lifecycle for P-SSP7 – MED4 interactions, the current results clearly demonstrate that the integrated production and preparation pipeline works to generate sufficient samples for in situ cryo-ET experiments.

**Figure 6.**
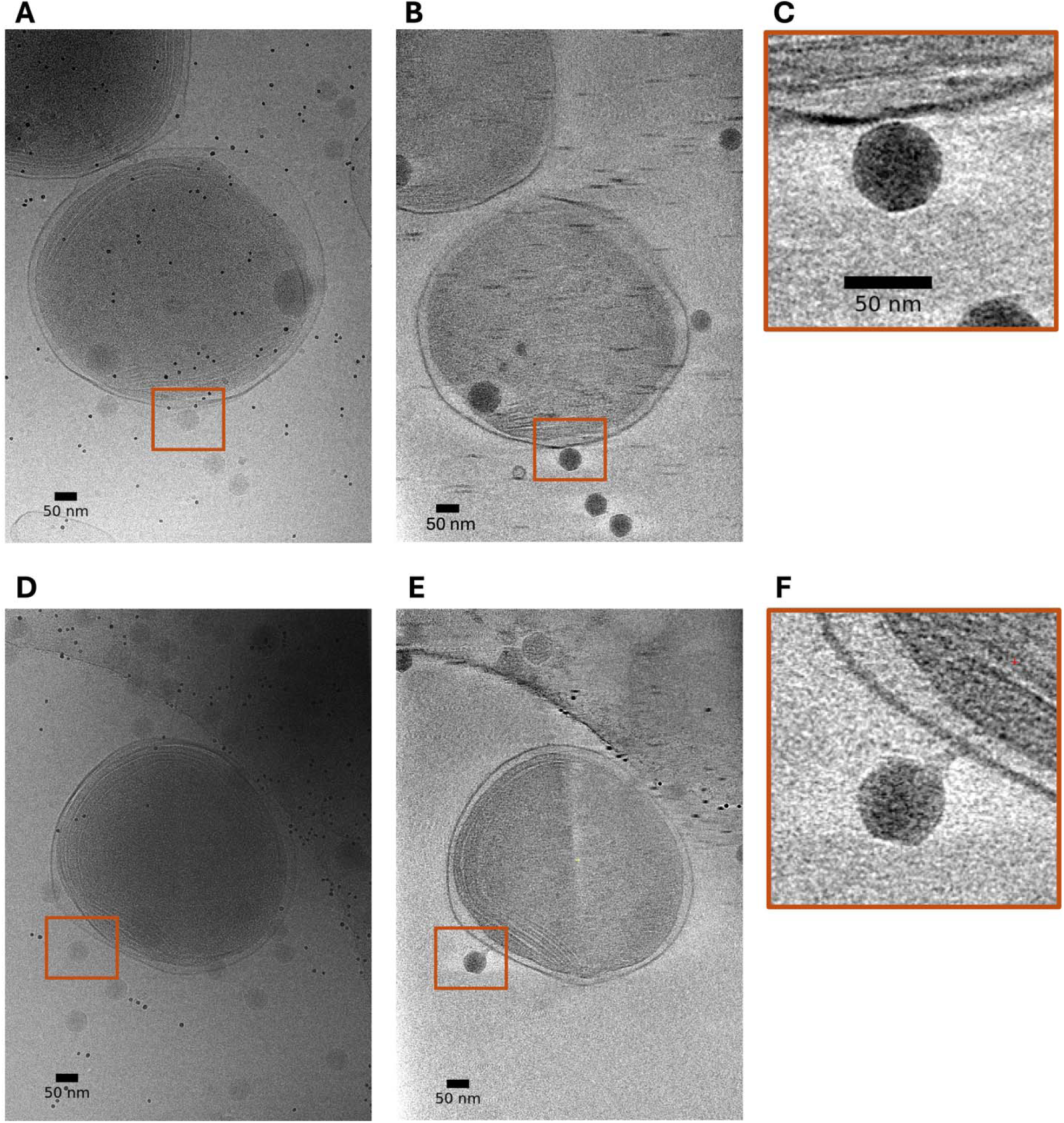
Cryo-ET visualization of 2 different states of host-phage interaction. A) Cryo-EM image at 0^0^ tilt of a PSSP-7 infected MED4 cell 15 minutes post infection. B) 45 nm thick slice through the reconstructed 3d volume (tomogram). C) Region in the brown boundary in (A) and (B) zoomed in to show the interaction between the phage and the host cell. D) Cryo-EM image at 0^0^ tilt of a PSSP-7 infected MED4 cell 45 minutes post infection. E) 45 nm thick slice through the reconstructed 3D volume (tomogram) showing a PSSP-7 phage in an intermediate stage of infection with the phage tail attached to the cell and orthogonal to the membrane. F) Region in the brown boundary in (D) and (E) zoomed in showing the tail firmly attached to the host membrane. Scale bar = 50 nm.

## 4. Discussion

This work demonstrates that systematic integration of cultivation, infection, and purification optimization can transform marine cyanophage studies from specialized, resource-intensive endeavor to routine research capabilities. The successful scale-up from small-volume maintenance cultures (10-100 mL) to laboratory production volumes up to 40 L represents a systematic technological advancement that makes high-quality marine cyanophage preparations routinely achievable across diverse research settings that meet their requirement of minimal biomass for multimodal experimental measurements. While previous structural studies of P-SSP7 have been accomplished, our integrated methodology provides comprehensive protocols that address the reproducibility, scalability, and multi-application challenges that have limited broader adoption of marine cyanophage structural biology.

More broadly, these advances bridge a critical gap in marine virus research, where marine cyanophages represent one of the most abundant biological entities in ocean ecosystems, with an estimated typical number of 10^7^ viral particles per mL in surface waters (Jacquet et al., 2010; Jameson et al., 2011), yet structural and multi-omics characterization has lagged far behind terrestrial phage systems. While high-resolution structures of marine cyanophage capsid proteins have been determined (Liu et al., 2010; Rao et al., 2021; Cai et al., 2023), comprehensive studies of portal-tail complexes remain constrained by sample quality limitations rather than technological limitations of cryo-EM itself (Passmore and Russo, 2016; Carragher et al., 2019; Nakane et al., 2020; Yip et al., 2020; Weissenberger et al., 2021). Such limitations have prevented the application of emerging visual proteomics approaches to marine virus systems. Suboptimal sample quality necessitates prohibitively extensive imaging campaigns to achieve adequate data for high-resolution reconstruction. Our methodology directly addresses the sample preparation bottleneck that has been identified as the primary limiting factor in achieving high-resolution viral structures (Passmore and Russo, 2016; Carragher et al., 2019; Weissenberger et al., 2021).

The most significant achievement of this methodology is the >60-fold improvement in cryo-EM data collection efficiency (**Table 2**), which transforms structural studies from requiring weeks of instrument access into single-day experiments that fit within typical facility allocation windows. Together, these advances reduce the barrier to entry for marine virus structural biology by an estimated order of magnitude in both time and cost, making comprehensive structural characterization accessible to the broader research community.

### Key Methodological Innovations

Our integrated approach addresses multiple bottlenecks simultaneously, distinguishing it from previous efforts that tackled individual challenges in isolation. While standardized protocols exist for laboratory and scaled production of bacteriophages for therapeutic applications (Luong et al., 2020; João et al., 2021; Kosznik-Kwaśnicka et al., 2024; Lapras et al., 2024; Luong et al., 2024; Wiebe et al., 2024), sustainable agriculture (Jo et al., 2024), viral metagenomics (Thurber et al., 2009), and laboratory stocks for common phages (*e.g.,* T4-types phages) (Bonilla et al., 2016), no comprehensive methodology has addressed the specific requirements of marine cyanophage production for high-resolution structural studies, requiring exceptional sample purity, particle concentrations, and stability optimized for cryo-EM applications. The systematic optimization of three key parameters (cultivation scalability, infection efficiency, and purification quality) enabled the dramatic improvements in sample quality necessary for routine structural studies.

Enhanced nutrient supplementation represents a critical factor controlling cyanophage infection success and productivity (Singh, 2012; Jassim and Limoges, 2013), yet its optimization is often overlooked. Our 2× nitrogen and phosphorus supplementation strategy directly addresses the intensive biosynthetic demands of phage replication, which begin within minutes of infection and require sustained extracellular nutrient acquisition (Stent and Maaløe, 1953; Waldbauer et al., 2019). Studies demonstrate that phosphorus limitation alone can reduce phage burst sizes by 73-85% and extend latent periods 2-fold or more (Cheng et al., 2019; Howard-Varona et al., 2024), while severe nutrient stress can lead to pseudolysogenic responses where infected cells fail to lyse (Wilson et al., 1996; Rihtman, 2016). This nutritional optimization contributed significantly to achieving the consistent high-titer phage production (3.1×10^12^ units/mL) across all production scales, providing the foundation for subsequent purification improvements.

Systematic MOI optimization represents a critical parameter that varies significantly across cyanophage studies, with values reported in the literature spanning from 10^-6^ to 10^3^ (Grasso et al., 2022), reflecting diverse experimental objectives and host-phage systems. For routine maintenance and scaled production, low MOI values (0.001-0.1) are typically employed to maximize phage amplification through sequential rounds of infection (Wilson et al., 1996; Zborowsky and Lindell, 2019; Laurenceau et al., 2021; Jo et al., 2024; Kosznik-Kwaśnicka et al., 2024; Wiebe et al., 2024), an approach that has proven efficient across various phage-host systems where MOI values of 0.01-0.1 consistently yield optimal titers (Jo et al., 2024; Luong et al., 2024; Jintasakul et al., 2025). However, MOI optimization remains system-specific, with factors including host growth characteristics, phage adsorption efficiency, and burst size influencing optimal values (Wiebe et al., 2024). Our systematic approach using freshly prepared lysate at optimized MOI values (<0.01 for maintenance, 0.005-0.01 for scaled production) aligns with established best practices while ensuring reproducible high-titer production suitable for structural applications. The combination of this MOI optimization with enhanced nutrient supplementation enabled the 100- to 1000-fold improvements in infectious titers observed between intermediate and large-scale preparations (**Table 1**).

The systematic adaptation strategy for local seawater sources addresses both economic and logistical challenges of large-scale production. Commercial seawater costs would exceed $3,400 plus shipping expenses for a typical 40 L production run, compared to minimal processing costs for local seawater. Beyond cost considerations, local sourcing provides operational flexibility by eliminating dependence on shipping schedules and supply chain constraints, enabling more responsive production planning.

The quality-driven optimization approach represents a departure from traditional viral production methods that prioritize yield over purity. Our systematic vessel scaling from PYREX bottles (**Supplementary Figure S1**) to PET carboys (**Supplementary Figure S2**) with corresponding aeration system modifications enabled this optimization while maintaining sterile conditions. As demonstrated by recent cryo-EM facility surveys (Alewijnse et al., 2017; Clare et al., 2017), sample quality rather than instrument capability has become the primary determinant of successful high-resolution structural determination. Our iterative optimization, guided by cryo-EM validation at each scale, exemplifies the integration of end-user requirements into bioprocess development that has been identified as critical for advancing structural biology applications (Carragher et al., 2019; Modafferi et al., 2025). The multi-stage approach, featuring enhanced protocols for debris removal, selective concentration, and two-step cesium chloride gradient purification, systematically addresses the complex contamination profile of cyanobacterial lysates (Wolf and Reichl, 2014; Tanir et al., 2021). This purification optimization, built upon the foundation of improved cultivation and infection protocols, ultimately enabled the >60-fold reduction in cryo-EM data collection time by achieving 66 particles per average field of view.

### Broader Applications and Future Directions

The methodology established here enables research applications that were previously less practical for marine virus systems. While developed specifically for the model marine cyanophage P-SSP7, the principles established—local seawater adaptation, optimized nutrient supplementation, and quality-driven purification workflows—should be transferable to other marine virus-host systems, providing a framework for process development across diverse viral families and ocean environments (Suttle, 2007; Gregory et al., 2019; Dominguez-Huerta et al., 2022).

The sample quality and throughput achieved create opportunities for applying emerging “visual proteomics” approaches (Huang et al., 2023; Strack, 2023) to marine cyanophage systems. The ability to routinely produce samples with >60 particles per field and >95% purity provides a foundation for more comprehensive structural characterization, potentially enabling systematic mapping of viral protein structures and virus-host interactions at high resolution. The reduction in data collection time from >60 days to ∼1 day makes previously resource-intensive studies more feasible, including single-particle reconstruction of portal-tail complexes and comparative structural studies across cyanophage families.

Future developments could further enhance accessibility and throughput. Process automation, particularly of PEG precipitation and dialysis procedures, could reduce labor requirements and improve reproducibility. The methodology’s success also opens possibilities for comparative structural studies across marine cyanophage families—studies that would have required weeks to months of instrument time using previous approaches but can now be completed within days—potentially revealing fundamental principles of marine virus evolution and host specificity mechanisms relevant to understanding ocean biogeochemical cycles (Rodriguez-Valera et al., 2009; Thompson et al., 2011; Flombaum et al., 2013).

## Acknowledgements

This work is partially supported by the NW-BRaVE for Biopreparedness project funded by the U. S. Department of Energy (DOE), Office of Science, Office of Biological and Environmental Research, under FWP 81832 and a BSSD Cryo-EM Operations project under FWP 74915. A portion of this research was performed on the project awards (10.46936/staf.proj.2023.61054/60012367 and 10.46936/expl.proj.2024.61521/60012928) from the Environmental Molecular Sciences Laboratory, a DOE Office of Science User Facility sponsored by the Biological and Environmental Research program under Contract No. DE-AC0576RL01830. Pacific Northwest National Laboratory is a multi-program national laboratory operated by Battelle for the DOE under Contract DE-AC05-76RL01830. We acknowledge PNNL-Sequim, including the Research and Facilities Staff that made access to Sequim Bay seawater possible for this effort. We particularly thank Scott Edmundson, who coordinated research associates and staff to collect and ship seawater for our cultivation scale-up campaigns, as well as the scientific divers who routinely clear the PNNL-Sequim lines, and facilities crew who maintain the seawater pumps and distribution pipes.

## Author contributions

Conceptualization – P.B and A.D.P. Methodology – P.B, A.D.P and N.C.S. Original draft writing – P.B. and A.D.P. Reviewing and editing – P.B, A.D.P, N.C.S and M.S.C and J.E.E. Supervision, project administration and funding – M.S.C and J.E.E

