## Supplementary Information for "Systematic Scale-Up and Enhanced Purification of Marine Cyanophage P-SSP7"

Description of **Figure S1. Custom-made intermediate-scale cultivation and air supply systems for *Prochlorococcus marinus* MED4 and infection experiments**

**(A) 1 L PYREX bottle-based cultivation setup with sterile air filtration and delivery system for intermediate-scale cultivation.** The system employs 1 L PYREX® Round Media Storage Bottles (#1395-1L, Corning Incorporated) that can be acid-washed, sterilized by autoclaving, and reused many times. The custom cap system allows delivery of filtration-sterilized air. The gentle bubbling system provides controlled CO₂ supply essential for photosynthetic growth while maintaining sterile conditions. Flow rates are maintained at 0.05-0.1 L/min to prevent excessive shear stress on fragile *Prochlorococcus* cells.

**(B) Custom cap system for intermediate-scale systems based on 1 or 2 L PYREX bottles.** Corning® polybutylene terephthalate (PBT) high-temperature caps (#1395-45DC) feature three pre-drilled holes accommodating 1/8" OD PTFE tubing for: (1) sterile air inlet with 0.2 μm filtration, (2) air outlet for pressure equilibration, and (3) sterile sampling port for culture monitoring. Acro™ 37 TF vent devices with 0.2 μm PTFE membrane (#4464, Cytiva) provide air sterilization, connected via silicone tubing to individual culture vessels. The cap design maintains airtight seals while enabling continuous gentle aeration required for sustained cultivation periods of 7-10 days during infection cycles. Flow rates are maintained at 0.05-0.1 L/min to provide controlled CO₂ supply essential for photosynthetic growth while preventing excessive shear stress on fragile *Prochlorococcus* cells.

**(C) Example of intermediate-scale cultivation array (1-6 L total volume) with live and active *Prochlorococcus marinus* MED4 cells in preparation for infection.** Six 1 L PYREX® Round Media Storage Bottles arranged for parallel processing, each equipped with custom aeration systems. Individual vessels contain 0.7-1 L of Pro99 medium prepared with locally adapted Salish Sea seawater and enhanced nutrient supplementation (2× nitrogen and phosphorus). The characteristic green coloration indicates healthy *Prochlorococcus* populations with active chlorophyll content suitable for efficient phage infection. The parallel configuration enables systematic optimization of infection parameters while maintaining consistent environmental conditions across replicates. Temperature control is maintained at 21°C with continuous illumination at 25 μmol quanta m⁻²⋅s⁻¹ during infection phases.

**(D) Same cultures post-infection with P-SSP7 phage featuring chlorolytic and near-completely lysed cultures of *Prochlorococcus marinus* MED4 host cells.** The characteristic lack of green coloration indicates lysed *Prochlorococcus* populations without active chlorophyll content as the result of phage infection. Left vessels show incomplete loss of chlorophyll resulting from slower progression of infection compared to three bottles in the center featuring very late-stage infection with extensive host cell lysis. This demonstrates the progression from healthy, green cultures to clear, lysed cultures containing released phage particles ready for harvesting and purification.

**(E) Example of intermediate-scale cultivation array (2-8 L total volume) with live and active *Prochlorococcus marinus* MED4 cells in preparation for infection.** Four 2 L PYREX® Round Media Storage Bottles (#1395-2L, Corning Incorporated) arranged for parallel processing, each equipped with custom aeration systems. Individual vessels contain 1.5-2 L of Pro99 medium prepared with locally adapted Salish Sea seawater and enhanced nutrient supplementation (2× nitrogen and phosphorus). This larger vessel format enables higher total phage yields while maintaining the established protocols for host cultivation, infection timing, and environmental control. The scaling from 1 L to 2 L vessels represents the systematic volume increase approach used throughout the optimization process.

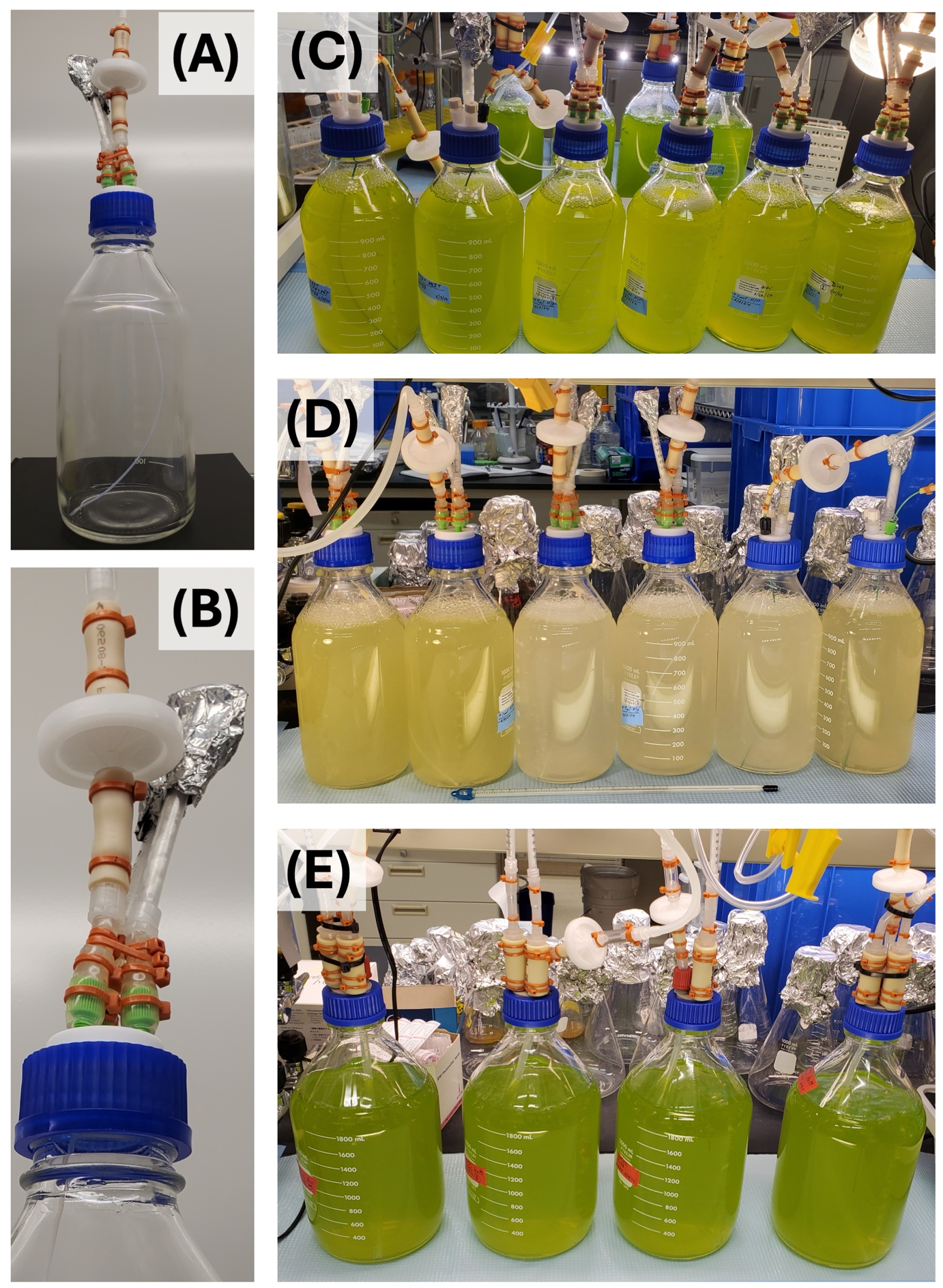

**Figure S1. Custom intermediate-scale cultivation and air supply systems for *Prochlorococcus marinus* MED4.**

Description of **Figure S2. Custom-made large-scale cultivation and air supply systems for *Prochlorococcus marinus* MED4 and infection experiments**

**(A) Food-grade PET carboy for large-scale cultivation.** Individual 12 L food-grade polyethylene terephthalate (PET) carboy designed for 10 L working volume with 2 L headspace to accommodate gas exchange during extended cultivation periods. The transparent PET material enables light penetration for photosynthetic growth and visual monitoring of infection progression while providing chemical compatibility with Pro99 marine medium and biological compatibility with *Prochlorococcus marinus* MED4. Carboys are sterilized using 70% ethanol rinse followed by sterile air drying, avoiding the need for autoclave sterilization as PET material is not resistant to steam-based sterilization.

**(B) Custom cap system for large-scale PET carboy systems.** Modified Jaece Industries Identi-plug™ Plastic Foam Stoppers (#L800-D) accommodate sterile aeration tubing while maintaining airtight seals. The foam stopper design allows for easy insertion of 1/8" OD PTFE tubing for air inlet and outlet connections while providing secure closure during extended cultivation periods. This system enables sterile air delivery and venting at optimized flow rates of 0.1-0.2 L/min per vessel, proportionally scaled for 10 L working volumes to maintain adequate CO₂ supply without excessive turbulence.

**(C) Sterile preparation of large-scale cultivation systems in biosafety cabinet.** Three PET carboys being prepared under sterile conditions with Pro99 medium containing locally adapted Salish Sea seawater and enhanced nutrient supplementation (2× nitrogen and phosphorus). Initially, 2 L of culture were transferred from intermediate-scale vessels and diluted 2-fold by adding fresh medium. As cells grow, additional medium is supplied to maintain cells in exponential phase until reaching 10 L of dense *P. marinus* culture. The characteristic green coloration indicates healthy, active *Prochlorococcus marinus* MED4 cultures ready for infection experiments. Sterile technique during setup prevents contamination during the extended cultivation and infection periods required for large-scale phage production.

**(D) Large-scale cultivation systems during operation in controlled environment chamber.** Multiple PET carboys positioned in a Conviron BDW series walk-in chamber providing precise temperature control at 21°C and continuous illumination at approximately 35 μmol quanta m⁻²⋅s⁻¹. The controlled environment ensures consistent growth conditions across all vessels during the critical infection phase. Individual aeration systems deliver filtered air through sterile tubing, maintaining optimal CO₂ levels while preventing cross-contamination between vessels. This setup represents the final scale-up phase capable of producing 40 L total cyanophage lysate volume for scaled production of high-quality phage enabling comprehensive structural studies.

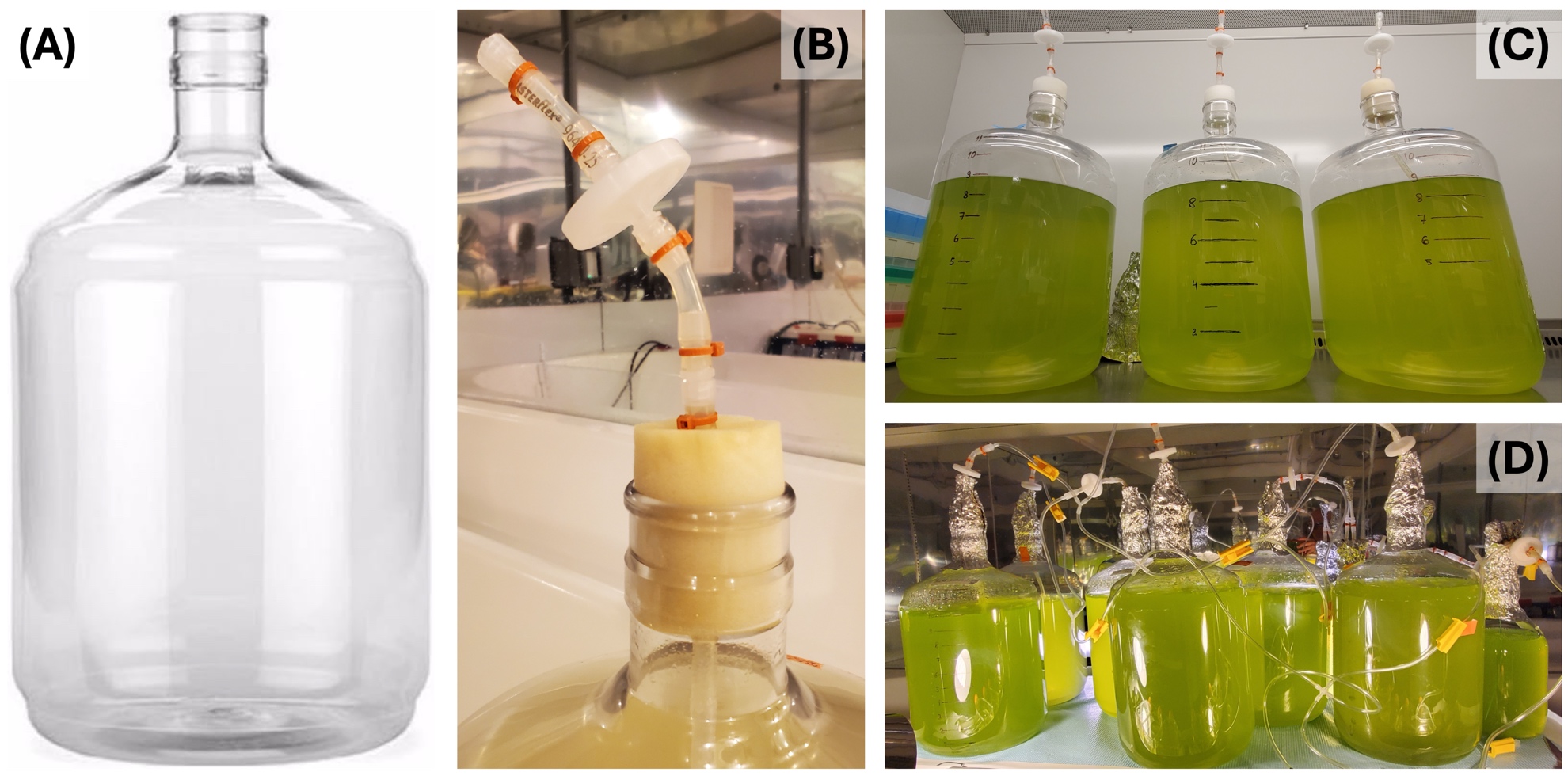
